# Novel single-cell preservation and RNA sequencing technology unlocks field studies for *Plasmodium* natural infections

**DOI:** 10.1101/2024.07.05.602255

**Authors:** Erin Sauve, Pieter Monsieurs, Pieter Guetens, Roberto Rudge de Moraes Barros, Anna Rosanas-Urgell

## Abstract

Single-cell RNA sequencing (scRNA-seq) is a powerful technology used to investigate cellular heterogeneity. When applied to unicellular eukaryotes such as *Plasmodium* parasites, scRNA-seq provides a single-cell resolution particularly valuable to study complex infections which are often comprised of mixed life stages and clones. Until now, the application of scRNA-seq has been mainly limited to *in vitro* and animal malaria models, despite known transcriptional differences as compared to circulating parasite populations. This is primarily due to the challenges of working with *Plasmodium* natural infections in endemic settings. We validated sample preparation methods and a novel single-cell RNA sequencing technology for the first time in *P. knowlesi* parasites which can be effectively implemented to analyze natural infections in low-resource settings. We recovered 22,345 *P. knowlesi* single-cell transcriptomes containing all asexual blood stages from 6 *in vitro* culture samples, with conditions mimicking natural infections, and generated the most extensive *P. knowlesi* single-cell dataset to date. All 6 samples produced reproducible circular UMAP projections with consistent cluster localization and high gene expression correlation, regardless of the sample preparation methods used. Biomarker expression and life stage annotation using the Malaria Cell Atlas *P. knowlesi* reference dataset further confirmed these results. In conclusion, the combination of adaptable sample preparation methods with novel preservation and scRNA-seq technology has the potential to fundamentally transform the study of natural infections. This approach unlocks the use of scRNA-seq in field studies which will lead to new insights into *Plasmodium* parasite biology.

**Importance:** Sequencing unicellular organisms, such as malaria parasites, at the single-cell level is important to understand the diversity present in cell populations. Until now, single-cell sequencing of malaria has been primarily limited to laboratory models. While these models are key to understanding biological processes, there are known differences between lab models and parasite populations circulating in natural human infections. This study presents sample preparation methods and a new single-cell RNA sequencing technology that enables sample collection from natural infections in low-resource settings. Using a mock natural infection, we validated this new single-cell RNA sequencing technology using marker genes with known expression patterns and a reference dataset from the Malaria Cell Atlas. We demonstrate that high-quality single-cell transcriptomes with consistent expression patterns can be recovered using various sample preparation methods, thereby unlocking single-cell sequencing for field studies and leading to additional insights into parasite biology in the future.

## Introduction

Single-cell RNA sequencing (scRNA-seq) offers a powerful approach to unravel biological complexity of unicellular eukaryotes, as it provides unprecedented resolution into cellular heterogeneity (1). This is particularly useful when investigating the expression profiles of single-celled organisms with complex life cycles and mixed life stages or mixed clones in parasite populations.

The application of scRNA-seq to *Plasmodium* parasites has been insightful to reconstruct cell trajectories and reveal transcriptional profiles associated with different stages in the intraerythrocytic developmental cycle across species (2–12). Studies focusing on *P. falciparum* (5–8) and *P. berghei* blood stages (9, 10) have begun to unravel gene expression signatures of sexual commitment and differentiation. Additional studies have described gene expression profiles of *Plasmodium* sporozoite development and liver stages in *P. berghei* (2, 13–16)*, P. falciparum* (17, 18), and *P. vivax* (12, 19–21). This effort has primarily been led by the Malaria Cell Atlas (MCA) which has the most comprehensive resource of publicly available *Plasmodium* single-cell data (2–4, 9, 17, 22).

Despite the vast potential of scRNA-seq in malaria research, its application has been largely restricted to parasites from *in vitro* cultures or animal models, with only one study analyzing four Malian natural infections being published to date (4). While laboratory strains are essential for studying *Plasmodium* biology, they are not necessarily representative of circulating parasite populations. Genome-wide transcriptomic studies have demonstrated inherent differences in gene expression between culture-adapted strains and natural infections (23–26). Moreover, natural infections are often complex with individuals frequently carrying multiple clones (27–31). Different clones within a single host can vary in their drug resistance, virulence, immune evasion or invasion strategies, providing insights into how parasites interact, compete, and evolve during an infection. The Dogga et al. scRNA-seq study analyzed four Malian natural infections revealing cell clusters with significantly lower gene expression levels as compared to laboratory strains, as well as differential expression between clones within the same host in genes related to host-parasite interactions (4). The capability to identify and characterize intra-host diversity using scRNA-seq will advance our understanding of the complexity of natural human infections and uncover critical aspects of gene expression in *Plasmodium* biology (32). For these reasons, it is important to develop and validate methods that extend single-cell transcriptomic studies to low-resource settings.

The challenges associated with applying omics techniques to study *Plasmodium* natural infections in humans, including the abundance of human genetic material, low parasite densities, and limitations of laboratory equipment at patient recruitment sites in low-resource endemic settings, extend beyond single-cell methodologies and are also commonly shared with bulk omics approaches. A combination of leukocyte depletion and parasite enrichment methods are indispensable to remove human genetic material and increase the concentration of parasites prior to RNA preservation (33–40). An additional challenge for scRNA-seq of human natural infections is the need to preserve cell integrity, which precludes the use of RNA preservation buffers which lyse cells. Furthermore, methods for scRNA-seq of natural infections need to be suited for transportation between field and laboratory sites, as sample collection and processing for scRNA-seq often occur at different locations.

Traditionally, the two main single-cell technologies used for *Plasmodium* are droplet- and well-based technologies. While these two technologies differ in their cell isolation methodology and sequencing resolution (41), they both require fresh cells and costly instruments in addition to equipment such as ultra-low temperature freezers, which are not often feasible in remote health centers. Recently, a new well-based technology has been developed with the advantage that it is instrument-free and provides integrated RNA preservation, making it compatible with sample collection in low-resource settings. HIVEs (Honeycomb Biotechnologies) are small devices that contain a microscopic well (pico-well) system designed to mimic a honeycomb which captures and preserves up to 60,000 single cells for the CLX version. Each pico-well contains a uniquely barcoded bead that captures mRNA using a poly-dT tail, similar to droplet sequencing. RNA preservation buffer is added to the HIVE before freezing to ensure RNA integrity and allowing the HIVEs to be stored for up to 9 months as per manufacturer’s instructions. Once frozen, HIVEs can be shipped on dry ice to a central laboratory for processing and library preparation, making it ideal for studies involving multiple collection sites or time points. During processing, HIVEs undergo bead collection, transcriptome recovery and amplification, and library preparation to create libraries that can be sequenced using standard short-read sequencing approaches.

In this study, we applied HIVE CLX single-cell sequencing technology for the first time to *Plasmodium* parasites and validated different parasite enrichment methods to make it deployable for the collection of cells from natural infections in low-resource endemic settings. Standardized parasite enrichment protocols combined with preservation and scRNA-seq technologies have the potential to revolutionize *Plasmodium* research by enabling scRNA-seq of natural infections.

## Results

### Processing *Plasmodium* natural infection samples to recover all erythrocytic life stages

To validate the application of HIVE technology for the analysis of *Plasmodium* natural infections, we tested different sample processing and loading methods, and assessed the single-cell RNA sequencing results. To create a mock natural infection sample with mixed parasite life stages, we diluted *in vitro*-cultured *P. knowlesi* A1-H.1 parasites to 0.8% parasitemia in whole blood.

Tested methods for sample processing include two combinations of leukodepletion and parasite enrichment techniques tailored to optimally isolate all life stages present in blood circulation and eliminate uninfected human cells from natural infections, which would otherwise dominate sequencing reads. A schematic representation of the protocols of the 6 samples is provided in Figure 1. For the PNyco method, which only requires a tube centrifuge, we used a Plasmodipur filter to remove human leukocytes, followed by a Nycodenz density gradient to enrich for parasite life stages. Here we found an enrichment to 22% parasitemia after the Nycodenz gradient by microscopy. In the second method, MACSPS, we used a MACS column to enrich for trophozoite and schizont life stages in the magnetic column based on the paramagnetic properties of hemozoin, similar to the MCA method (2). The first elution contains leukocytes, uninfected RBCs, and ring stages which have passed through without binding to the magnetic column. Subsequently, the first elution underwent further processing using a Plasmodipur filter to remove leukocytes, followed by a saponin lysis to remove uninfected RBCs, resulting in the isolation of the ring stage. In the second elution, the trophozoite and schizont stages bound to the magnetic column were released and recovered. The ring stages were then combined with trophozoites and schizonts recovered to obtain an enriched sample containing all life stages.

**Figure 1:**
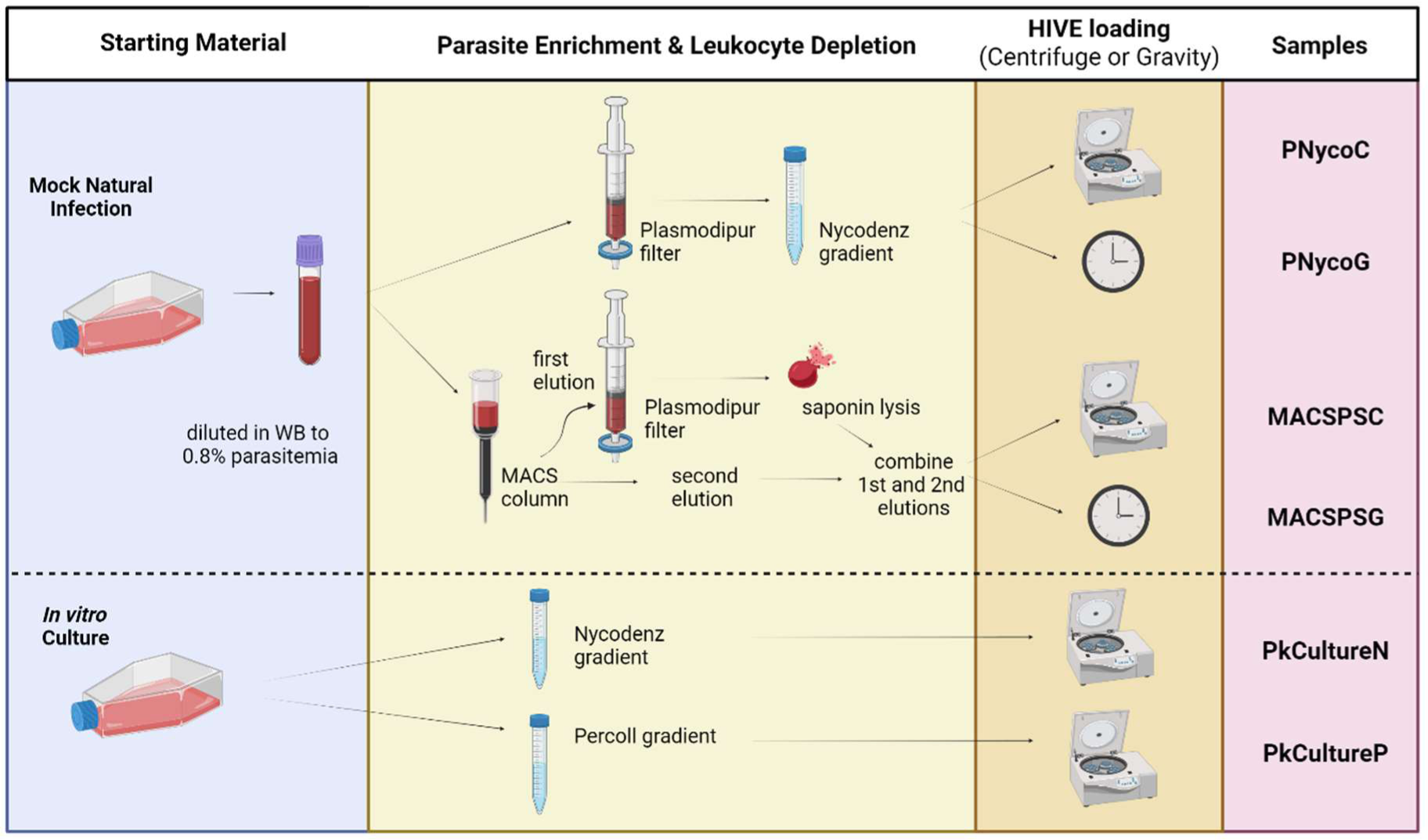
Experimental setup to test methods for processing *Plasmodium* natural infections and in vitro cultures. A mock natural infection sample was prepared by mixing *P. knowlesi* A1-H.1 *in vitro* culture with human whole blood. Two combinations of leukodepletion and parasite enrichment methods were tested (PNyco and MACSPS). In addition, we processed an *in vitro*-cultured *P. knowlesi* sample, with two different density gradients: Nycodenz (PkCultureN) and Percoll (PkCultureP). For each sample 60,000 cells were loaded into a HIVE by gravity (G) or centrifuge (C). Figure created using BioRender.com.

For the HIVE loading, we evaluated two different methods for loading enriched parasite into the HIVEs: one utilizing a plate centrifuge to facilitate the sinking of parasites into the pico-wells, and the other allowing parasites to sink by gravity, eliminating the need for a plate centrifuge if unavailable (C and G, respectively, Figure 1).

In addition, we also tested cultured *P. knowlesi* samples without mixing with whole blood. Parasite enrichment was conducted using two different density gradient centrifugation methods. Half of the cultured parasites were purified using a Percoll gradient (PkCultureP), typically employed to enrich samples for schizonts stages (40), while the other half underwent purification using a Nycodenz gradient (PkCultureN), commonly used for *P. knowlesi* (33), which recovered a broader range of parasite life stages compared to Percoll. Both methods led to a similar level of enrichment, around 50%, and samples were loaded using centrifugation.

### Validation of single-cell RNA sequencing using HIVE technology

A total number of 22,345 *P. knowlesi* parasite cells were recovered across 6 samples after applying a conservative filtering approach with thresholds of 300 genes and 600 transcripts per cell and doublet removal. The doublet detection and removal strategy applied to recover high-quality single cells is described in detail in the supplementary material (Figure S1 and Table S1). Cell recovery per sample ranges from 2,291 to 7,532 cells depending on enrichment and loading method (Figure 2) with a mean of 678 genes and 1,375 transcripts per cell (Figure S2 and Table S2).

**Figure 2:**
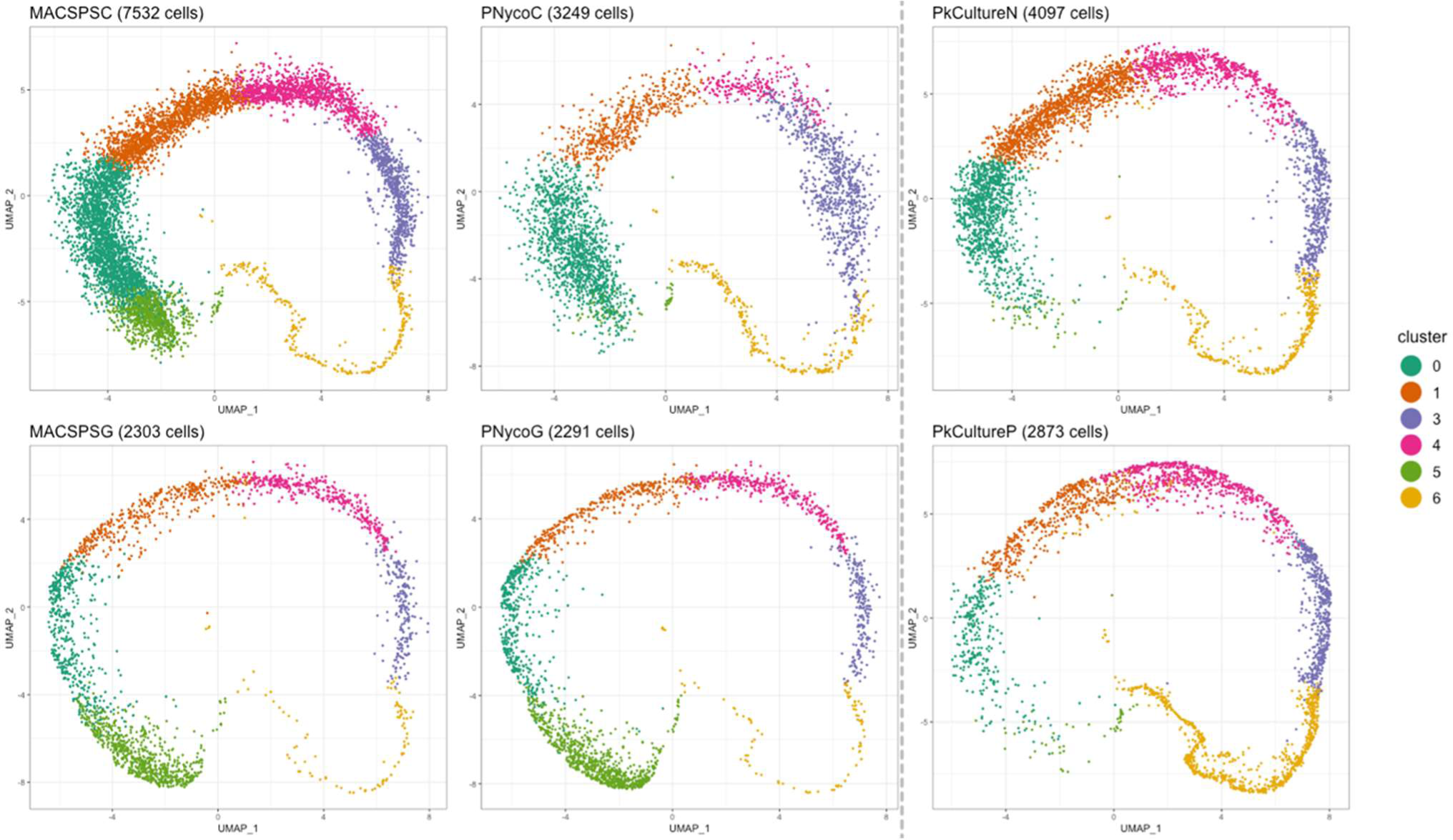
2D UMAP projections per sample by de novo clustering after doublet removal. Six clusters with comparable locations are observed in a ring formation across all sample UMAPs, with one cluster (cluster 2) being excluded from analysis during doublet removal.

We explored the single-cell transcriptome data of the 6 samples loaded in the HIVEs using uniform manifold approximation and projection (UMAP), which shows the cells in a circular pattern reproducible across the UMAPs of all samples and thus, processing methods (Figure 2). Moreover, *de novo* Louvain clustering on the PCA of the cell transcriptomes resulted in 7 clusters (cluster 2 was excluded during doublet removal), with consistent locations and partitioning of the clusters, despite differences in the number of cells recovered per cluster across the 6 samples. For example, as compared to the other samples, cells in cluster 5 are less abundant in PkCultureN, PkCultureP, and in the mock natural infection sample PNycoC (percentages lower than 2%), while cells in cluster 6 are most abundant in the PkCultureP sample processed with the Percoll density gradient (31%) as compared to the other samples (percentages lower than 10%).

Next, we investigated the transcriptomic variability using pairwise correlation plots, following a pseudo-bulk approach as described in the materials and methods section. We calculated the correlation of the gene expression values between different samples after summing per gene the number of reads over all cells (Figure 3). All samples show highly correlated gene expression profiles with Pearson’s correlation coefficients (R) greater than 0.88 (p<.001) in all pairwise comparisons, despite the use of the different sample types and processing methods. The strong correlation in expression profiles between the four mock natural infection samples (0.97) demonstrates that enrichment and loading methods did not affect the single-cell data generated. Correlation values of PkCultureN with mock samples ranges between 0.95 to 0.98; while correlation values of PkCultureP with mock samples is lower (0.88 and 0.90) likely reflective of the overall differences in cell abundance between clusters 5 and 6.

**Figure 3:**
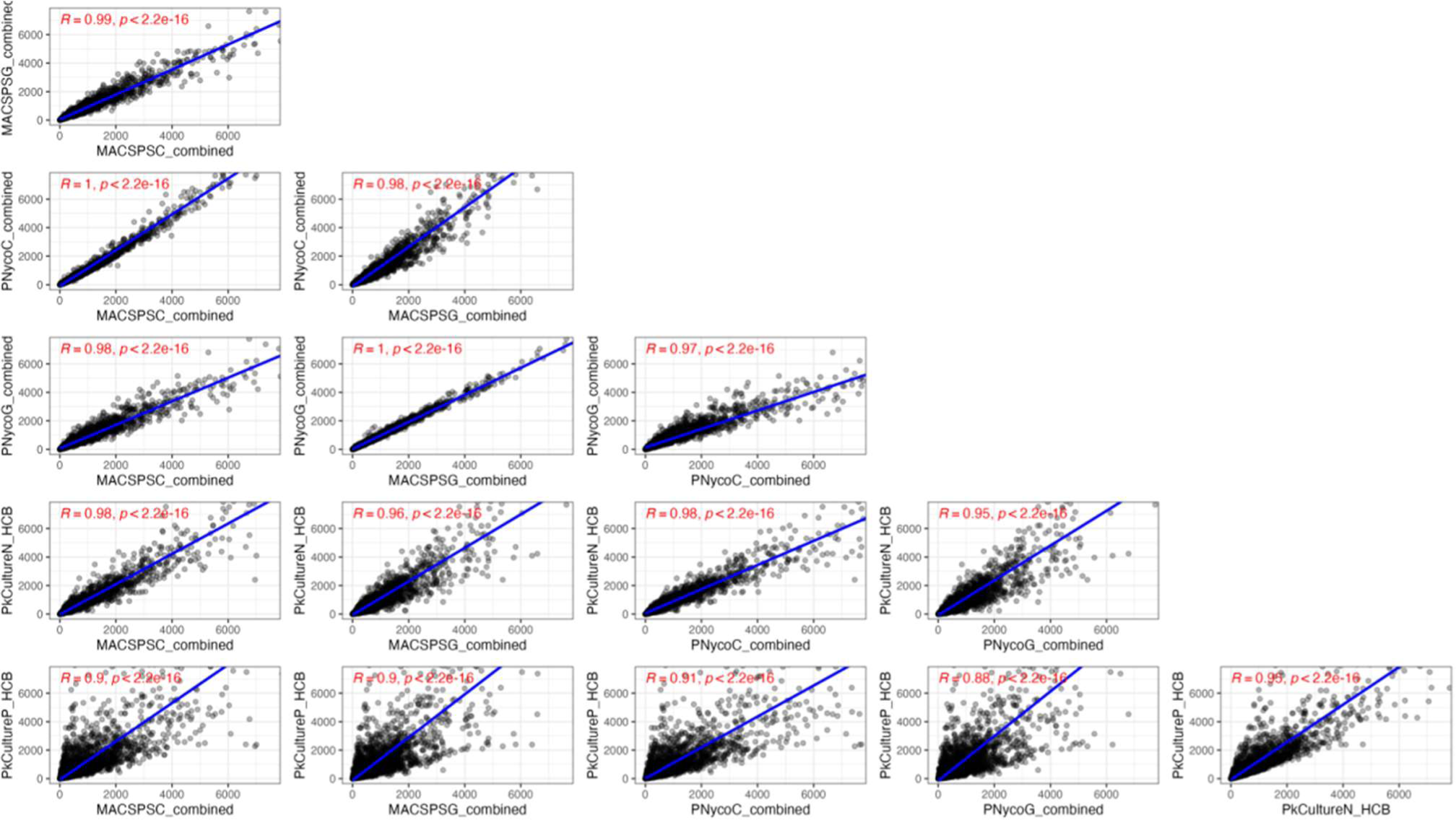
Pairwise correlation matrix of pseudo-bulk gene expression levels between sample processing and loading methods. Pearson correlation coefficients (R) and associated p-values are shown in red.

To examine whether HIVE sequencing technology is sensitive enough to detect gene expression at the single-cell level to study genes crucial for biological processes, we superimposed the expression levels of genes with known expression patterns to validate expression specificity (Figure 4). Tubulin (*pktubulin,* PKNH_0807700), a housekeeping gene which is consistently expressed throughout the lifecycle, is identified across all clusters in the UMAP (42, 43). The genes *merozoite surface protein 8, msp8* (PKNH_1031500), and *msp1* (PKNH_0728900) were used as markers of early and late blood stage asexual parasites, respectively (2). In the UMAP, the expression of *msp8* is highly enriched in cluster 5 and decreases in cluster 0 in contrast to expression of *msp1*, which begins in cluster 0 and increases clockwise with the highest level of expression in cluster 6.

**Figure 4:**
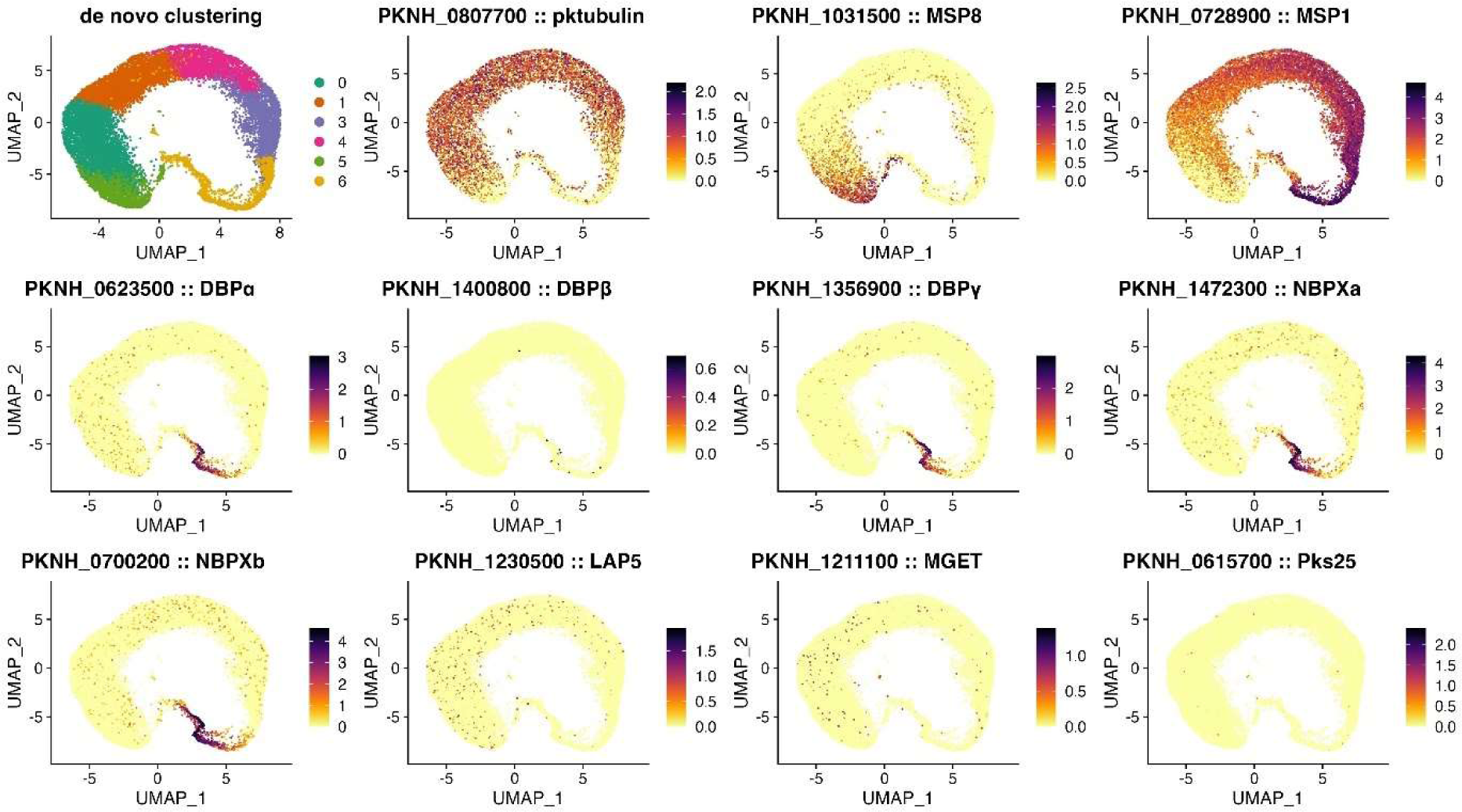
2D UMAP projections superimposed with gene expression levels (scaled and normalized counts) of marker genes show high and specific expression. First row (left to right): UMAP of all cells recovered with the *de novo* clustering annotation, *pktubulin* housekeeping gene with constant expression over the clusters, early (*msp8*) and late (*msp1*) asexual stage marker expression; Middle row (left to right): expression of genes involved in invasion (*PkDBPα, PkDBPβ, DBPγ and PkNBPXa*); Last row (left to right): *PkNBPXb* gene involved in invasion and gametocyte markers (*LAP5, MGET, Pks25*).

Genes involved in erythrocyte invasion, Duffy Binding Protein (DBP) genes alpha, beta, and gamma (*DBPα*, PKNH_0623500; *DBPβ*, PKNH_1400800; *DBPγ*, PKNH_1356900) and the homologues of the *P. vivax* Reticulocyte Binding Protein (*NBPXa*, PKNH_1472300 and *NBPXb*, PKNH_0700200), were projected due to their high-specific expression in the late schizont stage (25). These invasion genes, with the exception of *DBPβ*, are highly enriched in cluster 6. While *P. knowlesi* DBPα is a known ligand of DARC in human RBCs (44) and NBPXa has been identified as essential in human RBC invasion (45), the roles of DBPβ, DBPγ, and NBPXb remain unknown in human RBCs (46). We then examined the expression of known gametocyte marker genes, *LAP5* (PKNH_1230500), *MGET* (PKNH_1211100), and *ookinete surface protein Pks25* (PKNH_0615700) (11, 47). As the *P. knowlesi* A1-H.1 strain lost its ability to develop gametocytes when it was adapted to grow *in vitro* in human RBCs (33), we did not identify gametocyte gene markers in the UMAP, as expected.

Additionally, we used pseudotime analysis to further explore the gene expression of the *de novo* clustering and predict the developmental trajectory of the cells. Based on the mapping of life-stage specific marker genes *msp8* and *msp1* (Figure 4), clusters 5 and 6 were predefined as the start and end points of the pseudotime trajectory, respectively, which resulted in a single smooth trajectory without any branching (Figure 5A), indicating a single pathway of development. The circular-shape of the UMAP captures the cyclical nature of the asexual erythrocytic developmental trajectory as it maps the progression of gene expression and provides directionality, beginning after invasion in the ring stage, through the trophozoite stage, and ending in the schizont stage.

**Figure 5:**
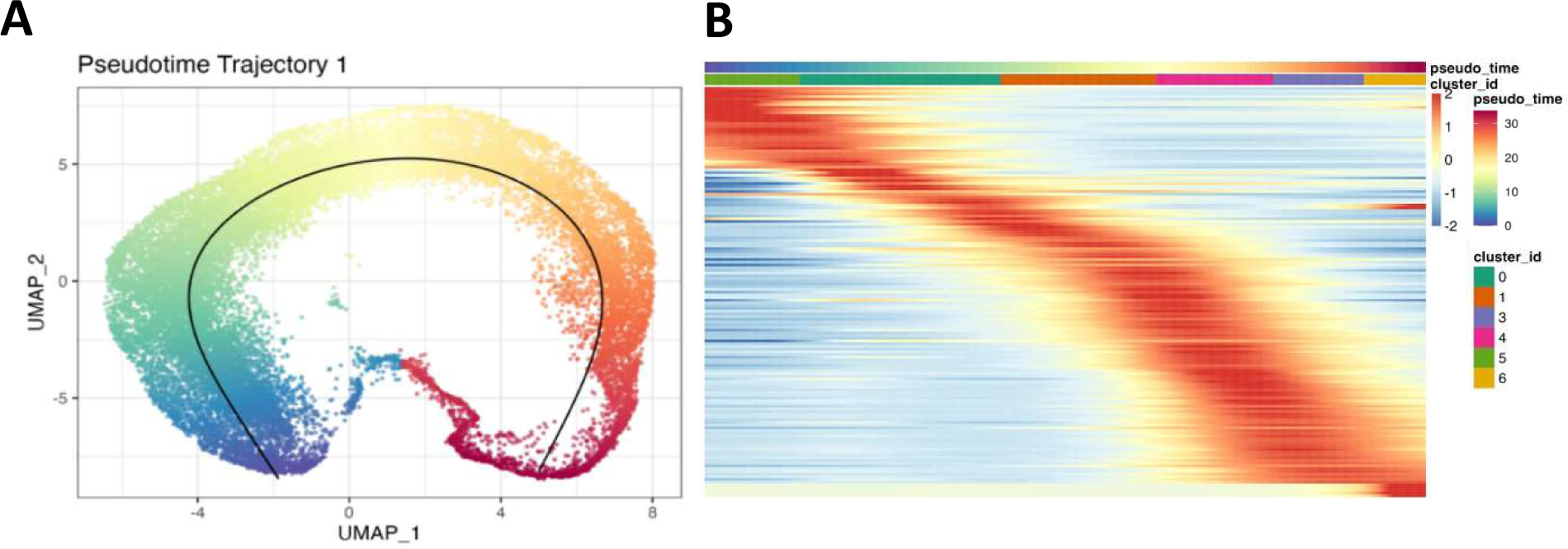
A) UMAP of the predicted developmental trajectory using pseudotime with the start and end points predefined as clusters 5 and 6, respectively. B) Heatmap showing the wave of gene expression (scaled and normalized counts) over the developmental trajectory by mapping the expression of the top 250 most variable genes (y-axis) by cell ordered by pseudotime and including the cluster annotation of each cell (x-axis).

The heatmap shows the top 250 genes (y-axis, Figure 5B) with the highest variable expression over the pseudotime corresponding with the identified *de novo* clustering (in the x-axis). The continuous biological transition is reflected by the wave-like changes in gene expression, which is in agreement with previous bulk and scRNA-seq studies (11, 25, 48), demonstrating that HIVE scRNA-seq can be used to study and map the dynamics of gene expression across the erythrocytic life cycle at the level of individual parasites.

### Life stage annotation and composition by enrichment method

Cells were assigned to the ring, trophozoite, or schizont life stages based on the scmap algorithm and the *P. knowlesi* MCA reference dataset (2). For each cell in our dataset, the scmap algorithm transfers the life stage annotation from the cell with the closest matching transcriptome in the MCA reference dataset. The annotated UMAP shows the recovery of all parasite life stages (Figure 6A) and that the location of the life stages corresponds well with the marker gene expression results in Figure 4. In addition, all life stages are recovered in all samples (Figure 6B), further confirming the reproducibility of the methods used. Out of the 22,345 cells in our dataset, only 2 cells could not be assigned (NA) to a specific life stage.

**Figure 6:**
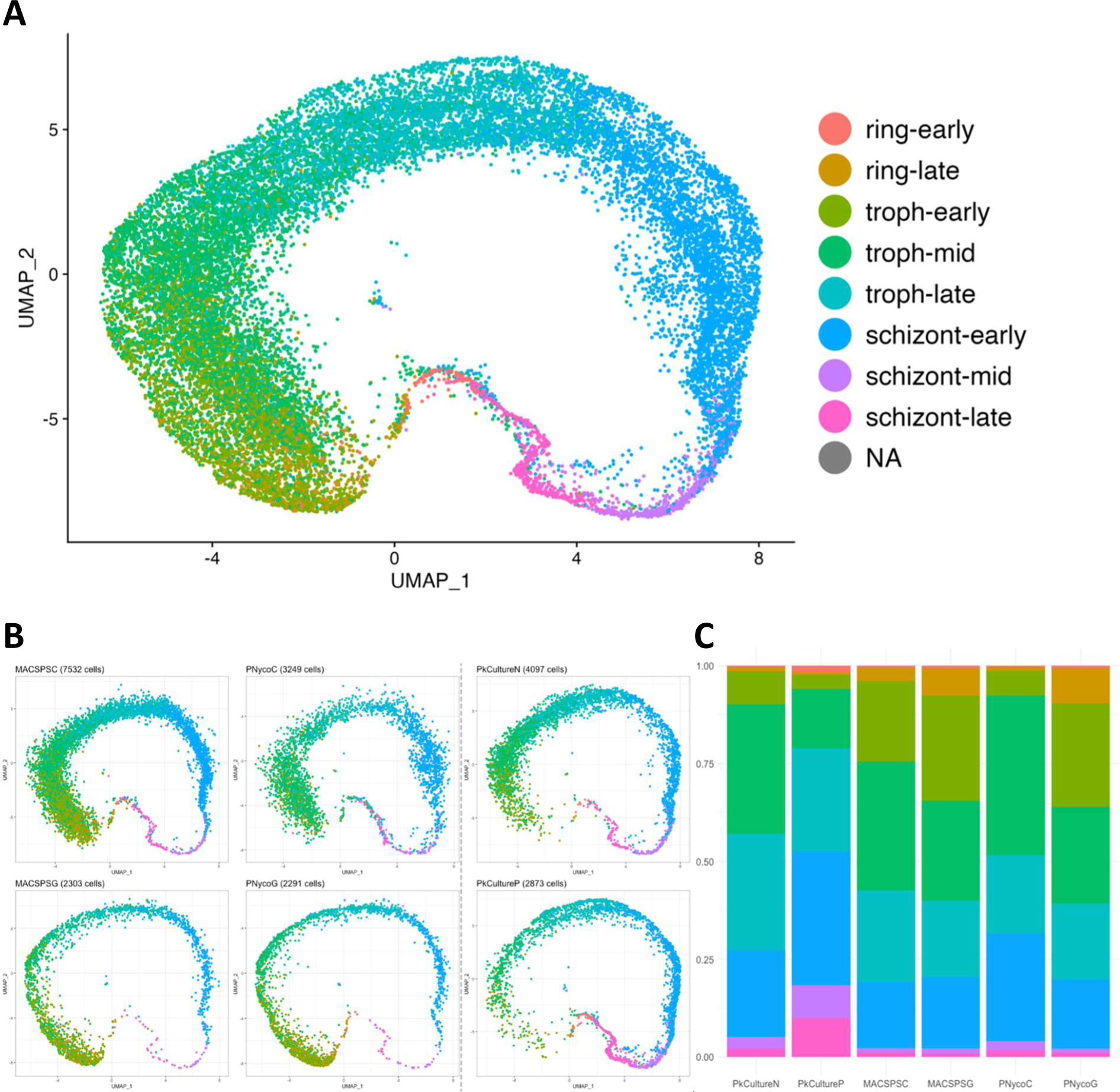
Life stage annotation of HIVE scRNA-seq dataset and life stage composition. A) 2D UMAP of the HIVE scRNA-seq dataset (22,345 cells) with the life stage annotation. Cells were assigned to ring (early/late), trophozoite (early/mid/late), and schizont (early/mid/late) life stages, using scmap and the *P. knowlesi* reference dataset. Only 2 cells could not be assigned a life stage (NA) B) 2D UMAPs showing the life stage annotation per sample C) Life stage composition recovered per sample as a percentage of the total number of cells recovered per method.

Subsequently, we compared cell and life stage recovery between sample processing and HIVE loading methods. Between the mock natural infection samples, the life stage composition remained remarkably stable between the processing (MACSPS vs PNyco) and loading methods (centrifuge vs gravity), although the number of cells recovered differed significantly (Figure 6C, Table 1). Mock natural infection samples loaded by gravity had a similar recovery of cells (2,303 cells vs 2,291, p>0.05) and life stage composition (8% vs 10% rings, 72% vs 71% trophozoites, 20% schizonts, p>0.05) regardless of processing method, MACSPS or PNyco, respectively (Table 1). Loading by gravity also showed a higher percentage recovery of ring stages as compared to loading by centrifuge (8% and 10% by gravity with MACSPS and PNyco vs 4% and 1% when loaded by centrifuge, respectively, both p<0.001); however, the total number of ring stage cells recovered by centrifuge is almost doubled (299 cells vs 176 by gravity with MACSPS), which is advantageous for the analysis.

**Table 1:** Cell recovery per sample indicating the absolute number of cells or by percentage of cell population among the 3 asexual erythrocytic life stages ring, trophozoite (troph), schizont. To calculate the percentages, the cell counts of the early, (mid), and late of each life stage were combined and divided by the total number of cells recovered for each sample.

| sample | Absolute numbers |  |  |  |  |  |  |  |  | Percentages |  |  |
| --- | --- | --- | --- | --- | --- | --- | --- | --- | --- | --- | --- | --- |
|  | ring |  | trophozoite |  |  | schizont |  |  | total | ring | troph | schizont |
|  | early | late | early | mid | late | early | mid | late |  |  |  |  |
| **PkCultureN** | 16 | 44 | 344 | 1356 | 1221 | 909 | 122 | 84 | **4096** | 1% | 71% | 27% |
| **PkCultureP** | 47 | 20 | 104 | 435 | 756 | 984 | 243 | 284 | **2873** | 2% | 45% | 53% |
| **MACSPSC** | 37 | 262 | 1550 | 2478 | 1757 | 1281 | 83 | 84 | **7532** | 4% | 77% | 19% |
| **MACSPSG** | 15 | 161 | 620 | 588 | 450 | 420 | 28 | 21 | **2303** | 8% | 72% | 20% |
| **PNycoC** | 9 | 34 | 205 | 1325 | 651 | 895 | 76 | 53 | **3248** | 1% | 67% | 32% |
| **PNycoG** | 14 | 207 | 606 | 566 | 444 | 406 | 23 | 25 | **2291** | 10% | 71% | 20% |

When looking at overall cell recovery, the MACSPSC method loaded by centrifuge recovered a 3-fold higher number of cells (7,532 cells vs 2,303 by gravity) and also the most cells of each life stage: ring, trophozoite, and schizont. The same comparison could unfortunately not be made for the PNyco method due to the unusually high percentage of removed doublets, which is most probably explained by an air bubble forming during the HIVE loading procedure. However, our analysis shows that while an air bubble can result in decreased cell recovery and a high number of doublets, the transcriptomes remaining after filtering are of high quality and the life stage composition is consistent, meaning that the cells can still be analyzed with the rest of the dataset.

In the PkCulture samples, The Nycodenz gradient (PkCultureN) recovered a significantly higher number of total cells (4,096 vs 2,873 cells, p<0.001) with a more even distribution over the developmental cycle than the Percoll gradient (PkCultureP). On the other hand, a Percoll gradient is highly efficient in enriching the sample for the schizont stage, with schizonts comprising 53% of the life stages recovered versus 27% with a Nycodenz gradient (p<0.001).

Across all samples, a smaller proportion of ring stages were recovered as compared to the percentage of rings present in the starting cultures (Figure S3). Before sample processing, the mock natural infection contained 33% rings, while our analysis identified 1% to 10% of the cells recovered as ring stages. Similarly, for the PkCulture samples, only 1 to 2% percent of cells recovered were rings, as opposed to the culture which contained 10% rings. This loss can partially be attributed to the difficulty of separating ring-infected RBCs from non-infected RBCs during sample processing, especially when using density gradients; however, there is an alternative explanation. To better understand the low recovery of rings, the number of genes and transcripts per cell were mapped for each cell ordered by their pseudotime (Figure 7). We found differences between the number of genes (Figure 7A) and transcripts (Figure 7B) detected per life stage, with rings having the lowest average number of genes and transcripts, trophozoites the highest, and schizonts having a similar number of transcripts despite expressing fewer genes. If we lower the thresholds to 100 genes and 250 transcripts per cell, we increase the recovery of rings from 866 to 2,287 cells, which indicates that when applying the conservative filtering thresholds of 300 genes and 600 transcripts per cell uniformly across the life stages, we are potentially filtering out high-quality rings that were successfully sequenced.

**Figure 7:**
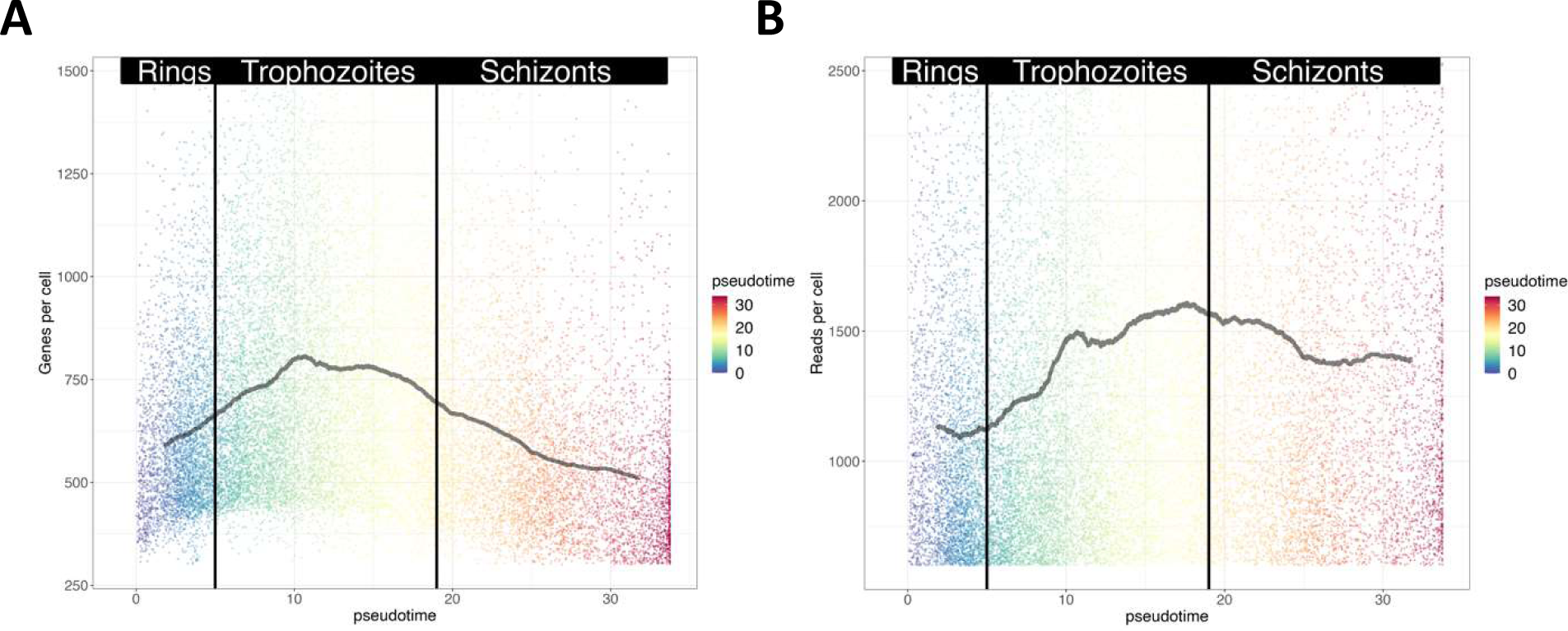
The number of genes (A) and number of transcripts (reads) (B) mapped per cell with the cells ordered by pseudotime to show the differences in number of genes and transcripts expressed over the developmental trajectory. While the transition between life stages is gradual, vertical lines have been superimposed to help distinguish between life stages.

## Discussion

Our study provides the first application and validation of HIVE CLX scRNA-seq technology to unicellular organisms and demonstrates the quality of the single-cell dataset generated by showing high reproducibility between samples and benchmarking our results with the publicly available MCA dataset. HIVE technology is unique in that it enables natural infection samples to be collected in low-resource settings as it is instrument-free and integrates RNA preservation, which allows sample collection to be separated from library preparation and sequencing.

We tested different sample processing and enrichment methods to isolate parasites from natural infections and cultured parasites as input for scRNA-seq using the HIVE technology. The methods tested successfully recovered all asexual erythrocytic life stages present with slight differences in life stage recovery between the methods. These differences can be leveraged to study different aspects of parasite biology. While erythrocytic sexual stages were not recovered since the *P. knowlesi* A1-H.1 strain lost its ability to develop gametocytes, we expect the MACS and density gradient protocols to also be suitable (49–51). For the cultured samples, a Percoll density gradient enriches more prominently for the schizont life stage, while a Nycodenz gradient performs a more balanced enrichment across the trophozoite and schizont stages. Therefore, Percoll can be best used to study egress and invasion, while Nycodenz is better suited to study the developmental trajectory. Additionally, both Percoll and Nycodenz density gradient percentages can be adjusted to enrich for different species and life stages (33, 39). The ring stage is notoriously difficult to enrich from natural infections due to their similarity in morphology to uninfected RBC and a lack of hemozoin (52). During parasite maturation inside the RBC, parasites digest hemoglobin, which lowers the density of the infected RBC (iRBC) and an increases hemozoin formation (the result of heme detoxification) (53–55). These changes in the iRBC enable the separation of trophozoite and schizont stages from uninfected RBCs using a density gradient or based on their paramagnetic hemozoin with a MACS column (35–37). However, newly infected iRBCs (ring stage) have not yet consumed enough hemoglobin to alter the density of the RBC nor produced enough hemozoin to be captured by a MACS column.

The MACSPS method, which combines a MACS column with Plasmodipur leukodepletion and saponin RBC lysis, was designed to maximize parasite enrichment (particularly of the ring stage) and remove uninfected RBCs from natural infection samples (2). When using this method and loading HIVEs by gravity, 8% of the total cells recovered were ring stages. This was highly comparable to the 10% rings stages obtained using the PNycoG method, but lower than expected after ring enrichment. Furthermore, when loading the MACSPS method by centrifuge, only 4% of the cells recovered were rings, despite the absolute number of ring cells recovered being twice as high. This discrepancy is likely due to the higher percentage of the less-dense late trophozoite and schizont stages settling into the wells with the help of the force from the centrifuge, thereby making the rings a smaller percentage of the total number of cells recovered.

Since all loading methods produce high-quality and comparable cell transcriptomes, loading by gravity initially seems ideal as it results in the highest percentage of ring recovery and may be the only option in some settings. However, the power of single-cell analysis relies on the number of cells, and thus to truly maximize cell recovery we recommend loading HIVEs using a centrifuge when available. Nevertheless, the high correlation between methods provides optimal versatility allowing for methods to be readily adapted to equipment availability. For example, a study with multiple collection sites could choose different processing and loading methods for each site and still analyze the samples together.

We generated, after filtering, scRNA-seq data of 22,345 *P. knowlesi* cells, more than five times the number of *P. knowlesi* cells published to date. Our *de novo* clustering reveals consistent clustering, with the same number of clusters in the same locations in all samples, which corresponds well with expected specific, localized biomarker expression and life stage annotation from the MCA. There is a clear shift from the egress of the schizont stage to the ring stage after invasion and continuing through the developmental trajectory, indicating biological significance of the clustering. While some overlap is seen in the life stage annotation, this is due to the use of an external reference to map life stages. Additionally, we show that there are no artifacts introduced into our dataset from the sample processing methods or preservation. The six samples included in the study were analyzed without the use of data integration methods (e.g. the anchor-based integration in Seurat (56) or Harmony (57)) often needed to remove batch effects between samples, demonstrating the high reproducibility of our methods, even when the PkCulture samples and mock natural infection samples were loaded at different times.

One advantage of the HIVE CLX sample capture is that the loading capacity is 60,000 cells. This is notably higher than the limitation of the number of wells used in SmartSeq2 sequencing or the maximum recommended loading of 16,000 cells with 10x Genomics droplet sequencing. The increased loading capacity of the HIVE CLX is particularly important when working with sample processing methods where populations are mixtures of iRBCs and RBCs. While enrichment techniques are helpful, they rarely lead to a pure population of iRBCs. From the 60,000 cells loaded per HIVE, we recovered up to 7,500 cells when loading by centrifuge. This represents 45% of the expected cell recovery reported by Honeycomb Biotechnologies. To maximize cell recovery for natural infection samples, future studies should sequence at a deeper level, ensuring a minimal number of 25,000 reads per cell for each sample, compared to the average depth of 16,925 reads per cell in this study.

To ensure that only high-quality cells were analyzed, we used stringent filtering thresholds of 300 genes and 600 transcripts per cell, as per the settings suggested in the Honeycomb vignette for the analysis of HIVE scRNA-seq data with Seurat. These thresholds are conservative compared to other *Plasmodium* scRNA-seq studies of erythrocytic life stages, which defined thresholds at 150 genes per cell for *P. knowlesi* or even lower for *P. falciparum* (2, 6). While conservative thresholds ensure data quality, they may impact the number of cells recovered, especially ring stages. Ring stages are characterized by lower transcription levels as compared to trophozoite and schizont stages (2, 6), making them less likely to pass conservative filtering and be included in the analysis. To avoid excessive filtering, thresholds could be adapted to each life stage. Optimizing filtering thresholds for ring stage recovery would be better explored in studies predominantly featuring ring stage cells. Adaptive filtering optimization is particularly important for natural infection samples, which often have low parasite densities and a high proportion of rings in circulation.

HIVE scRNA-seq provides high quality sequencing data that will lead to new insights into biological processes of *Plasmodium* and has similar potential for other unicellular organisms. The value of future studies investigating natural infections, especially for species where little is known such as *P. vivax,* and mixed-clone infections, should not be underestimated and should become widespread practice to understand *Plasmodium* biology now that it is accessible and adaptable to all settings.

## Methods

### Culture of *Plasmodium knowlesi*

*P. knowlesi* A1-H.1 strain parasites were maintained between 0.5 and 10% parasitemia in culture medium consisting of RPMI-1640 (Westburg Life Sciences) supplemented with 0.32 g/L sodium bicarbonate, 5.0 g/L Albumax II (Invitrogen, cat: 11021-029), 2.0 g/L D-glucose, 25mM HEPES, 0.05g/L hypoxanthine, 0.005 g/L gentamicin, 2mM L-glutamine with 10% (v/v) heat-inactivated horse serum (Invitrogen) at 3% hematocrit (HCT). Parasites were continuously cultured in O+ blood and incubated at 37°C under atmospheric conditions of 90% N2, 5% O2, 5% CO2. Parasitemia was determined using thin smear slides stained with 10% Giemsa stain for 10 minutes.

### HIVE Sample Processing

#### *P. knowlesi* culture enrichment with Nycodenz and Percoll density gradients

A *P. knowlesi* infected red blood cell (iRBC) pellet of 5% parasitemia was diluted to 50% HCT. 0.75mL of diluted culture was loaded onto 5mL of either 70% Nycodenz (N) (Serumwerk Bernburg, 18003) or 60% Percoll (P) (CAT NO) density gradient and centrifuged at 900g for 12 minutes (brake 0) for Nycodenz or 2400rpm for 10 minutes (brake 0) for Percoll, to separate iRBCs from non-iRBCs. The interfaces of the gradients were washed in 10mL medium without serum. The enriched iRBC pellets were resuspended in 2mL PBS-BSA buffer (PBS with 0.5% BSA).

#### *P. knowlesi* mock natural infection sample and parasite enrichment

A *P. knowlesi* iRBC pellet was diluted in 4mL whole blood to 0.8% parasitemia. The infected blood sample was centrifuged at 2400rpm for 10 minutes and buffy coat was removed. The sample was split into 2 aliquots and processed through two different methods (MACSPS and PNyco) for leukocyte depletion and parasite enrichment.

### MACSPS

One aliquot was processed using a MACS LD column (Miltenyi Biotec), a Plasmodipur filter (Europroxima), and saponin lysis (Sigma, cat: 47036). First, a MACS column, attached to a magnet, was pre-wet with PBS-BSA pre-warmed to 37°C. The iRBC pellet was diluted to 20% HCT and loaded onto the column. The column was washed three times with 1mL of PBS-BSA which recovered elution 1 containing mostly ring stages (elution 1). The column was removed from the magnet, washed twice with 3mL PBS-BSA and pelleted to elute the hemozoin-containing trophozoite and schizont stages (elution 2). Elution 1 was passed through a pre-wet Plasmodipur filter to deplete leukocytes, pelleted, and RBCs were lysed with 2.5V 0.15% (w/v) cold saponin (to remove non-iRBC) by pipetting 10x and pelleted for 5 minutes at 2400rpm. The pellet was washed twice with PBS, combined with the elution 2 pellet, and diluted in 2mL PBS-BSA.

### PNyco

The second aliquot was processed using a Plasmodipur filter and 70% Nycodenz density gradient prepared in PBS (PNyco). The aliquot was diluted to 20% HCT in PBS, passed through a pre-wet filter, and washed with 10mL PBS. The flow-through was pelleted, diluted to 50% HCT, and loaded onto a 70% Nycodenz gradient. This was centrifuged at 900g for 12 minutes (brake 0), the interface was washed in PBS, and 10μL of pellet was diluted in 2mL PBS-BSA.

### HIVE Sample Capture

Resuspended enriched iRBC pellets were further diluted in PBS-BSA to a concentration of 60,000 cells/mL using a Scepter 3.0 cell counter (Merck). 1mL of cell suspension was loaded into each HIVE to settle into the pico-wells via gravity or using a centrifuge according to modified manufacturer’s directions. The incubation time for the HIVE loading was increased from 30 minutes to 1 hour at room temperature (gravity) and from 3 to 5 minutes (centrifuge). HIVEs were stored at -20°C until library preparation to mimic using a -20°C freezer in the field.

### HIVE Library Preparation and Sequencing

Library preparation was conducted following manufacturer’s directions with 10 extra PCR cycles added to the whole transcriptome amplification (WTA). All samples were sequenced by on a NovaSeq6000 (Broad Institute, USA) according to the manufacturer’s recommendations. Sequencing data has been deposited in the NCBI sequencing read archive.

### Analysis

BeeNet v1.1 was used to map sequencing reads to an updated PKNH reference (downloaded from PlasmoDB version 62 (58)) with default parameters and an expected number of cells of 60,000. As 3’UTR annotations are absent for most *P. knowlesi* genes, the annotations of the 3′UTR were added in a systematic manner to enhance the accuracy of read assignments to the annotated transcript: for each protein-coding gene, the 3′UTR region was annotated by adding 2500 base pairs downstream of the stop codon. In cases where the newly designated 3′UTR overlapped with other genomic elements, the 3’UTR was truncated to avoid overlap. The resulting TCM and RCM files were read into R (version 4.2.1) and converted into a Seurat object using Seurat 4.2 (59). The single-cell data from 6 samples were merged into one Seurat object. Cells with less than 600 reads per cell or less than 300 expressed genes per cell were filtered out. Doublets were removed using scrublet (version 0.2.3, (60)) Preliminary analysis (PCA analysis and clustering analysis) before excluding potential doublets showed that one cluster was significantly enriched with scrublet-predicted doublets, and as such the entire cluster was also removed from the single-cell dataset. Count data were normalized using regularized negative binomial regression (SCTransform). After principal component analysis, the Uniform Manifold Approximation and Projection (UMAP) dimensional reduction technique was applied for visualization of the single-cell transcriptomes using the first 20 principal components (61). Clustering of cells was performed on the PCA output using a resolution of 0.25.

The single-cell data were aggregated into pseudo-bulk expression profiles by summing counts per gene across all cells within each sample. Pairwise correlations between samples were then calculated using the Pearson correlation coefficient via the *cor* function in R. Visualization of the correlation matrix was performed using the *ggplot2* package providing an overview of the similarity in gene expression profiles between samples.

Annotation of the life stages as described for *P. knowlesi* in the Malaria Cell Atlas (MCA) (2) was downloaded from https://zenodo.org/records/2843883 (pk10xIDC dataset), where the stage_pred column in the phenotypic metadata was used as life stage predictor. *P. knowlesi* single-cell transcriptomes obtained from the MCA database were projected on our dataset using the scmap-cell (version 1.18.0, (62)) implementation, and the corresponding annotation of the MCA cells was transferred to our dataset. Pseudotime analysis was performed using the Slingshot package (version 2.4.0, (63)) in R.

Statistical analysis was performed using R (version 4.3.3) with differences in cell recovery and life stage composition being determined using the Pearson chi-squared test.

## Acknowledgments

We would like to thank Dr. Ellen Knuepfer for donating the *P. knowlesi* A1-H.1 parasite strain which was used in this study. This work was funded by the Department of Economy, Science and Innovation in Flanders (SOFI to ARU, 2020-2024) and (jPPP to ARU and RMB, 2022). The computational resources and services used in this work were provided by the VSC (Flemish Supercomputer Center), funded by the Research Foundation - Flanders (FWO) and the Flemish Government. ES received support from an FWO PhD-SB Fellowship from the Research Foundation Flanders (1SC5522N). The funders had no role in study design, data collection, analysis, and interpretation, manuscript preparation, or the decision to submit the work for publication.

## Supplementary Material

**Figure S1:**
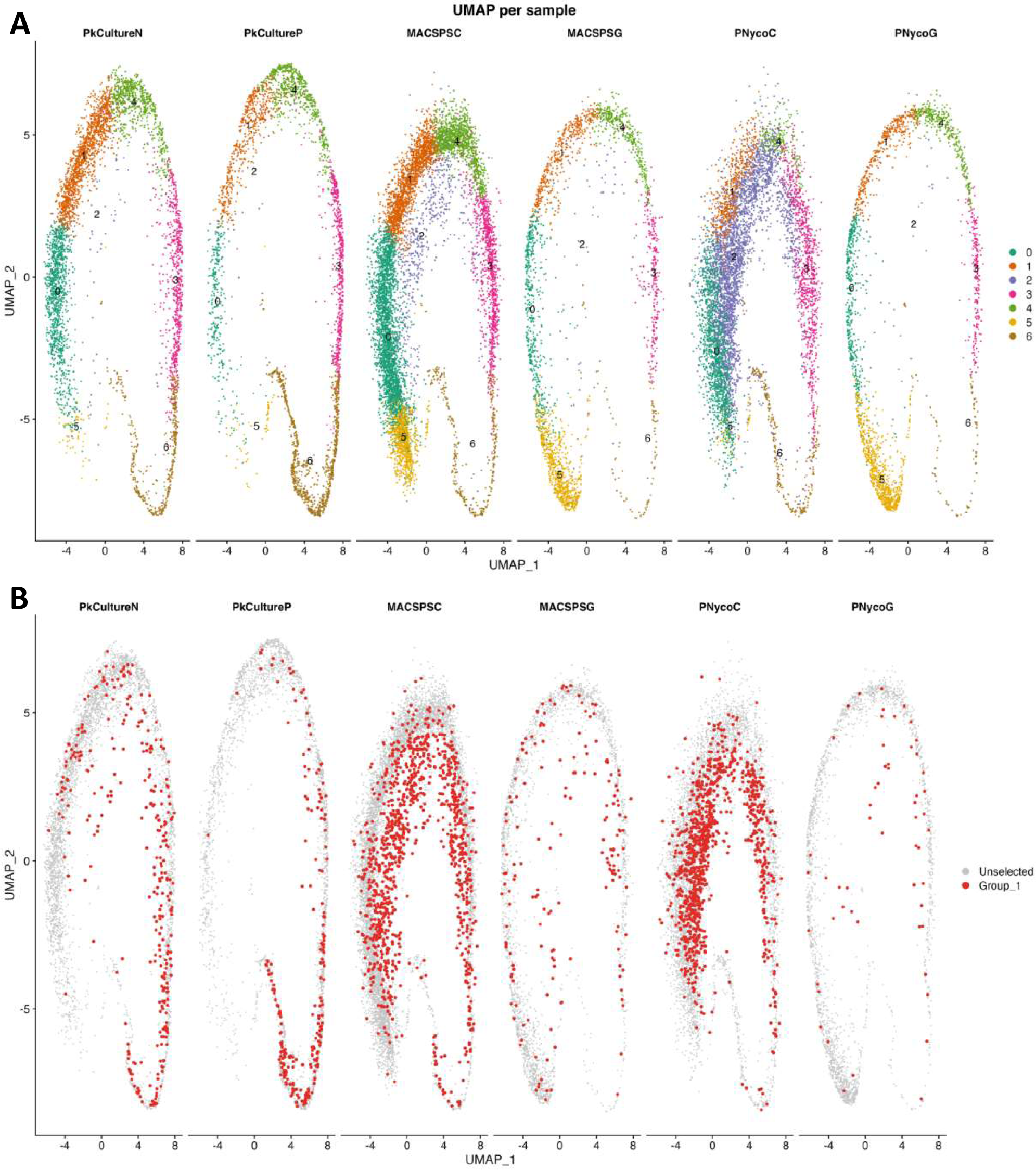
Visualization of doublet detection. A) showing the de novo clusters from the UMAP per sample before doublet removal, with cluster 2 which is eventually removed, B) UMAP showing the cells detected by Scrublet as doublets in red. These cells overlap well with cluster 2 in Part A of the figure seen in purple. Scrublet (v 0.2.3) was used to detect doublets, wells that contained multiple cells, in our dataset. The number of doublets detected and removed, between 3-14% for each sample (Table S1), fell within the reported ranges based on the cells loaded and recovered, except for sample PNycoC where more doublets were identified as compared to other methods, over 52%. This discrepancy in the number of doublets can likely be attributed to an air bubble discovered after centrifuging the PNycoC sample which created a dry patch on the membrane and thus concentrated the cells onto a smaller surface area than intended, leading to multiple cells in a single well. In addition to removing all identified doublets, cluster 2 was removed in its entirety as more than 50% of cells in the cluster were identified as doublets in half of the samples (Table S1). In addition, cells in cluster 2 mapped towards the middle of UMAP indicating these wells were likely showing an average expressions of more than one cell. In an effort to be conservative and to lower the likelihood that doublets, or multiple cells masking as a single-cell, remained in our dataset, cluster 2 was removed entirely from the analysis.

**Table S1.** Table describing doublet removal using Scrublet. The second column contains the number of cells before any doublets were removed. The third column, Scrublet doublet prediction, shows the number of cells predicted as doublets by Scrublet and the percentage of doublets as compared to the initial number of cells (second column). The fourth column, Cluster 2 (“doublet cluster”), describes the number and percentage of the cells belonging to cluster 2. The fifth column, Overlap scrublet with cluster 2, show the number and percentage of cluster 2 cells predicted as doublets, and the sixth column the number of doublet cells not overlapping with cluster 2. The seventh column shows the total number of doublets and the percentage of total cells that will be removed, and the last column shows the number of cells retained after doublet removal.

| Sample | # of cells (before) | Scrublet doublet prediction |  | Cluster 2 ("doublet cluster") |  | Overlap scrublet with cluster 2 |  | scrublet doublets not overlapping with cluster 2 | total doublets removed |  | # of cells (after) |
| --- | --- | --- | --- | --- | --- | --- | --- | --- | --- | --- | --- |
|  |  | # of cells | % of total cells | # of cells | % of total cells | # of cells | % of doublet cluster | # of cells | # of cells | % of total cells |  |
| MACSPSC | 8790 | 1050 | 12% | 605 | 7% | 397 | 66% | 653 | 1258 | 14% | 7532 |
| MACSPSG | 2512 | 174 | 7% | 81 | 3% | 46 | 57% | 128 | 209 | 8% | 2303 |
| PNycoC | 6787 | 990 | 15% | 3315 | 49% | 767 | 23% | 223 | 3538 | 52% | 3249 |
| PNycoG | 2360 | 55 | 2% | 33 | 1% | 19 | 58% | 36 | 69 | 3% | 2291 |
| PkCultureN | 4472 | 307 | 7% | 114 | 3% | 46 | 40% | 261 | 375 | 8% | 4097 |
| PkCultureP | 3134 | 232 | 7% | 35 | 1% | 6 | 17% | 226 | 261 | 8% | 2873 |
| Total | 28055 | 2808 | 10% | 4183 | 15% | 1281 | 31% | 1527 | 5710 | 20% | 22345 |

**Figure S2:**
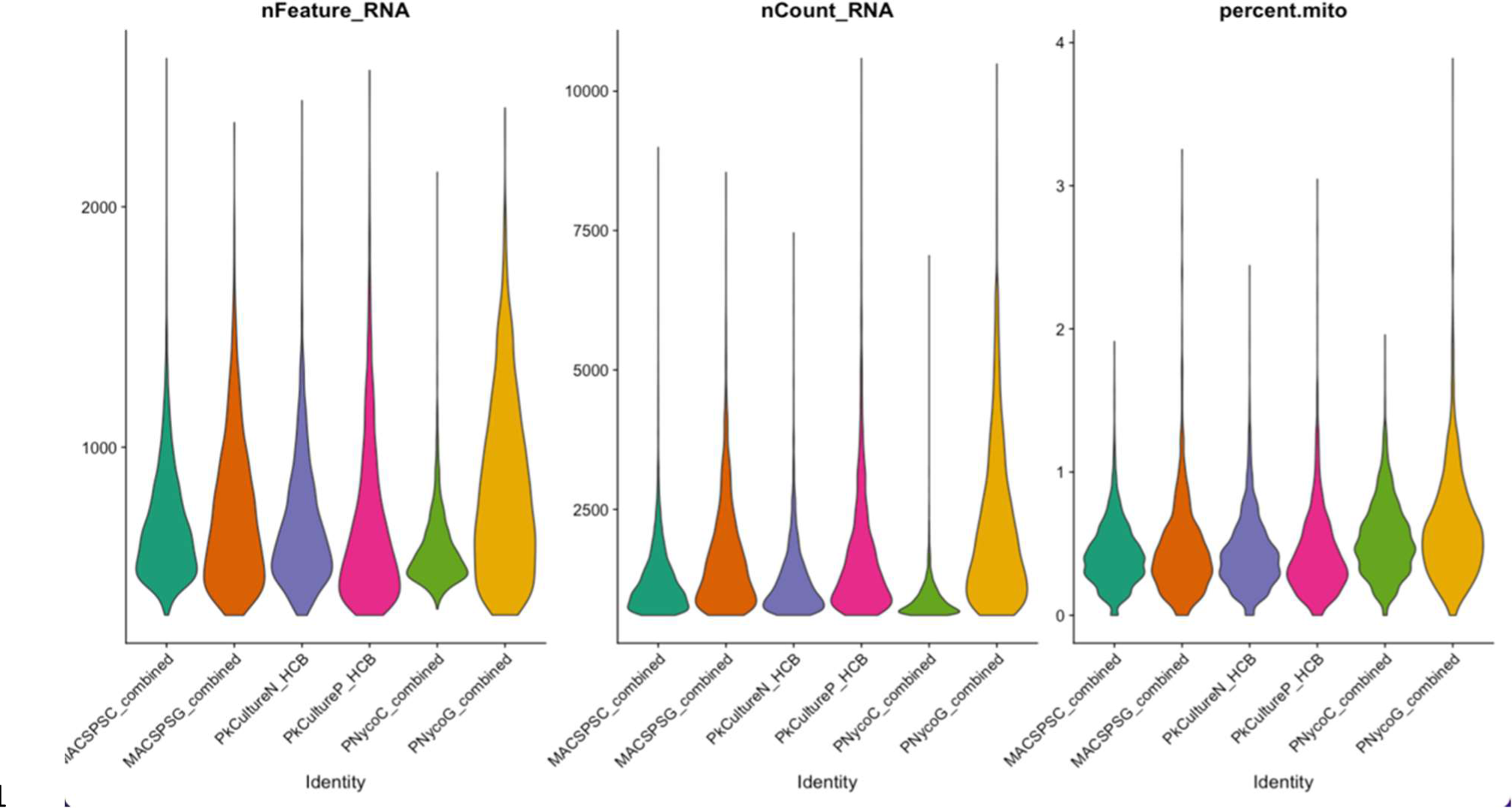
Quality Control of the scRNA-seq data. A) the number of genes per cell per sample, B) the number of reads (transcripts) per cell per sample and C) percent mitochondrial reads per sample

**Table S2:**
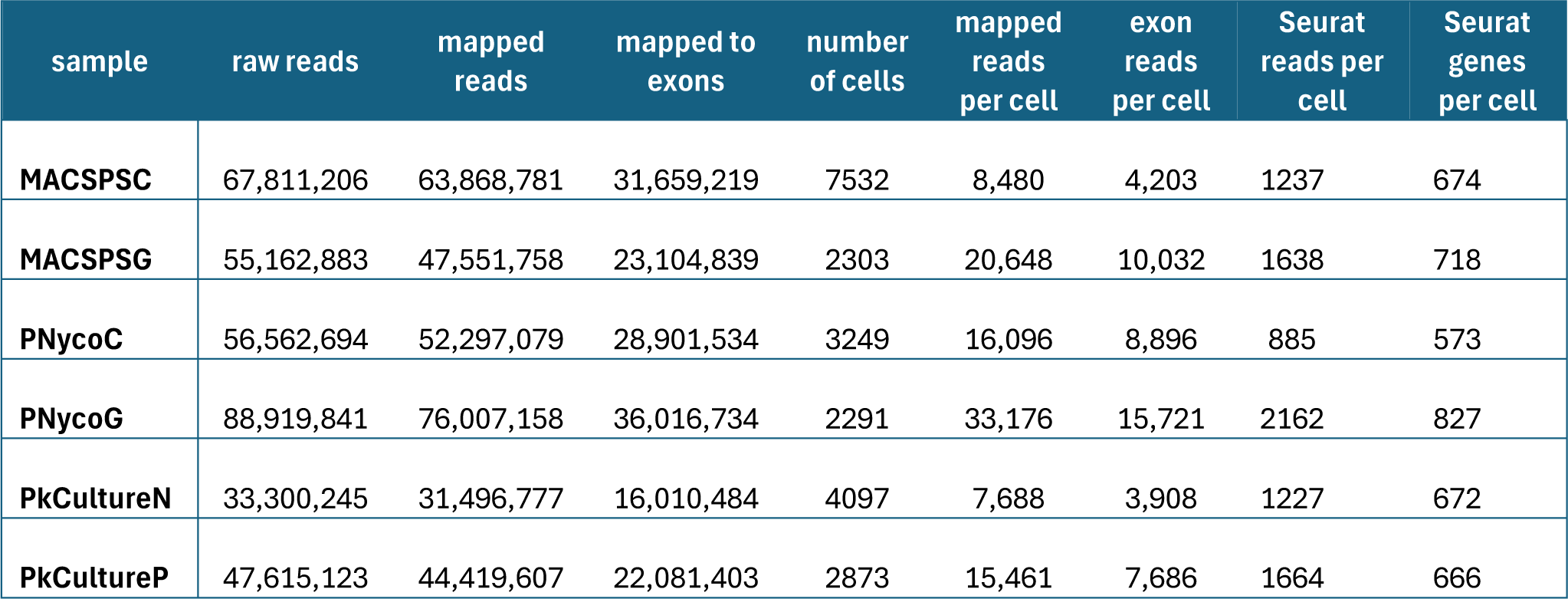
Sequencing statistics showing the raw reads, reads mapped to the *P. knowlesi* reference, the number of reads mapped to the exons, cells recovered after filtering, average number of reads (or transcripts) per cell and average number of genes per cell.

**Table S3:** Cell recovery per cluster per sample after filtering and doublet removal by absolute counts and percentages.

| sample | Absolute numbers |  |  |  |  |  |  | Percentages |  |  |  |  |  |
| --- | --- | --- | --- | --- | --- | --- | --- | --- | --- | --- | --- | --- | --- |
|  | 0 | 1 | 3 | 4 | 5 | 6 | total | 0 | 1 | 3 | 4 | 5 | 6 |
| **MACSPSC** | 2636 | 1676 | 766 | 1184 | 958 | 312 | **7532** | 35% | 22% | 10% | 16% | 13% | 4% |
| **MACSPSG** | 463 | 431 | 221 | 363 | 734 | 91 | **2303** | 20% | 19% | 10% | 16% | 32% | 4% |
| **PNycoC** | 1370 | 526 | 793 | 216 | 71 | 273 | **3249** | 42% | 16% | 24% | 7% | 2% | 8% |
| **PNycoG** | 442 | 373 | 210 | 365 | 809 | 92 | **2291** | 19% | 16% | 9% | 16% | 35% | 4% |
| **PkCultureN** | 1123 | 1315 | 519 | 669 | 60 | 411 | **4097** | 27% | 32% | 13% | 16% | 1% | 10% |
| **PkCultureP** | 292 | 509 | 526 | 609 | 55 | 882 | **2873** | 10% | 18% | 18% | 21% | 2% | 31% |

**Figure S3:**
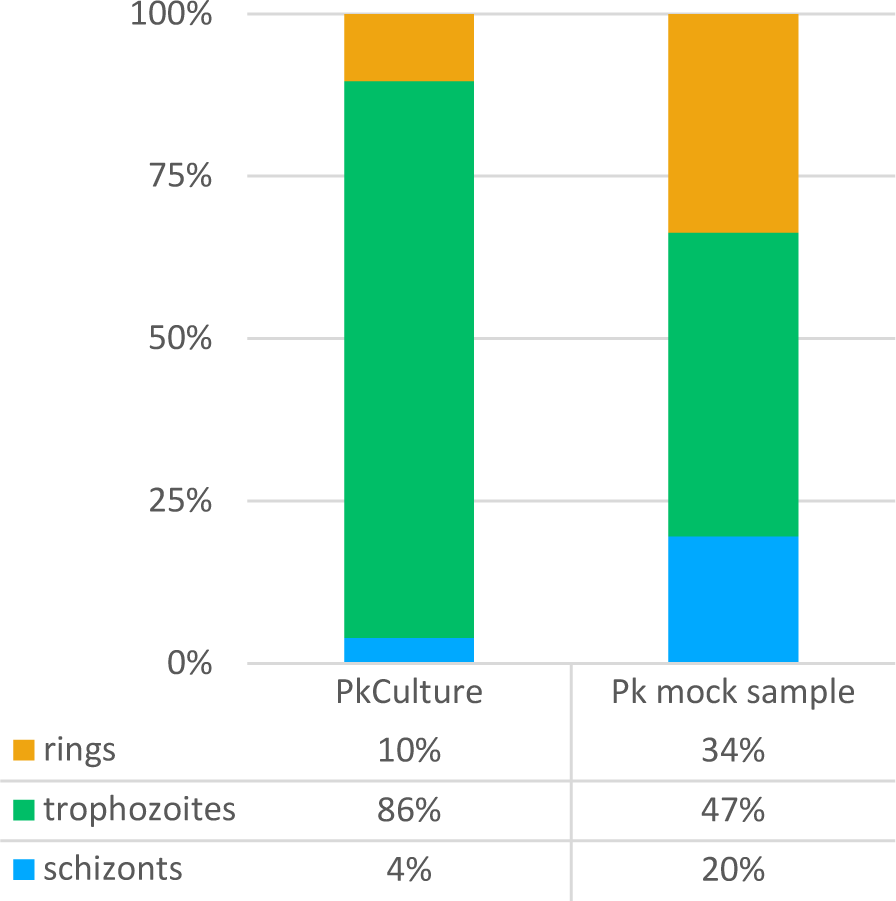
Life stage composition of the initial starting samples, PkCulture and Pk mock natural infection sample, by microscopy.

